# Error reduction in leukemia machine learning classification with conformal prediction

**DOI:** 10.1101/2024.12.11.627902

**Authors:** Mariya Lysenkova Wiklander, Dave Zachariah, Olga Krali, Jessica Nordlund

## Abstract

**Purpose:** Recent advances in machine learning (ML) have led to the development of classifiers that predict molecular subtypes of acute lymphoblastic leukemia (ALL) using RNA sequencing (RNA-seq) data. While these models have shown promising results, they often lack robust performance guarantees. The aim of this study was three-fold: to quantify the uncertainty of these classifiers; to provide prediction sets that control the false negative rate (FNR); and to perform implicit reduction by transforming incorrect predictions into uncertain predictions.

**Methods:** Conformal prediction is a distribution-agnostic framework for generating statistically calibrated prediction sets whose size reflects model uncertainty. In this study, we applied an extension called conformal risk control to ALLIUM, an RNA-seq ALL subtype classifier. Leveraging RNA-seq data from 1042 patient samples taken at diagnosis, we developed a multi-class conformal predictor, ALLCoP, which generates statistically guaranteed FNR-controlled prediction sets.

**Results:** ALLCoP was able to create prediction sets with specified FNR tolerances ranging from 7.5-30%. In a validation cohort, ALLCoP successfully reduced the FNR of the ALLIUM classifier from 8.95% to 3.5%. For cases whose subtype was not previously known, the use of ALLCoP was able to reduce the occurrence of empty predictions from 37% to 17%. Notably, up to 34% of the multiple-class prediction sets included the PAX5alt subtype, suggesting that increased prediction set size may reflect secondary aberrations and biological complexity, contributing to classifier uncertainty. Finally, ALLCoP was validated on two additional RNA-seq ALL subtype classifiers, ALLSorts and ALLCatchR.

**Conclusion:** Our results highlight the potential of conformal prediction in enhancing the use of oncological RNA-seq subtyping classifiers and also in uncovering additional molecular aberrations of potential clinical importance.

## INTRODUCTION

In the last decade, there has been an explosion in the number of diagnostic machine learning (ML) models developed for precision oncology, with the promise of delivering increasingly accurate diagnostics and personalized treatment ^1,2^. Acute lymphoblastic leukemia (ALL), the most common cancer in children and a highly heterogeneous disease, has seen the development of numerous classifiers linking transcriptomic footprints to subtype-defining chromosomal aberrations ^3–7^. ALL subtypes hold prognostic value, aid in monitoring disease progression, and help determine treatment intensity ^8,9^. Prior to the introduction of NGS-based diagnostics, ALL patients were subtyped at diagnosis using techniques such as G-banding, fluorescence in situ hybridization (FISH), and reverse transcription polymerase chain reaction (RT-PCR). Today, whole genome sequencing (WGS) and whole transcriptome sequencing (WTS) are emerging as alternative methods to determine subtype without a priori knowledge of the underlying genomic aberrations^10^. However, some patients remain unclassified even with the use of WTS analysis pipelines^11^, making ML subtyping classifiers a useful alternative. Yet, despite increasing efforts to implement data for clinical decision making in hematology^12^, numerous challenges remain to the deployment of diagnostic ML models, ranging from insufficient regulatory frameworks for artificial intelligence (AI) to the poor clinical applicability of such models ^13,14^.

Among these challenges is the quantification of reliability and uncertainty in ML classification models. Out of the box, classifiers typically output a naive*point prediction*, a top-scoring class or *k* top-scoring classes with no metric of uncertainty, while the underlying probabilistic scores for each class are not calibrated to empirical probability and are therefore not interpretable as confidence scores. Further, traditional ML models are evaluated with population-based metrics, which do not give useful indications of uncertainty for individual patients, and leave no explanation for classifier errors when they occur ^15^.

*Conformal prediction* (CP), a distribution- and model-agnostic framework, addresses this problem by replacing point predictions with *prediction sets* containing all true classes at a user-specified probability, using an independent calibration dataset to determine softmax thresholds for inclusion of classes in these sets ^16–19^. In addition to providing a mathematically proven, statistically guaranteed prediction set, CP aids the human interpretability of ML outputs: a CP set containing a single class shows that the classifier is highly certain about the prediction, while a larger set shows a higher degree of uncertainty, and an empty set indicates that the model does not recognize the input, suggesting an out-of-distribution (OOD) instance. Cresswell et al show that the use of CP sets, with size inherently quantifying uncertainty, can improve human decision making compared to models simply outputting the *k* top-scoring classes^20^. Finally, CP can be used for implicit error reduction: it is possible to select a lower error rate for the classifier (increased accuracy) at the expense of larger, less certain prediction sets (decreased precision). This is of particular utility for classifiers that have been trained on a small sample size or have underrepresented classes in the training data, such as ALL subtyping classifiers with high uncertainty in predicting the rarer subtypes^21^, or other under-trained models where more data becomes available over time.

To date, medical applications of CP remain few^22^. A handful of studies have shown CP applied to drug discovery^23^, disease course prediction in multiple sclerosis^24^, lung tissue microscopy^25^, as well as oncology, including prostate^26^ and breast cancer^27^ classification. A 2006 study applied CP to a Support Vector Machine trained to predict five ALL and three AML subtypes using microarray-generated gene expression data up to a 95% confidence level^21^. However, to date, there have been no applications of CP to RNA-seq classifiers for leukemia, or any other cancer.

In this study, we applied CP to the RNA-seq ALL subtype classifier ALLIUM^3^. A conformal predictor, ALLCoP, was calibrated and cross-validated using ALLIUM predictions across 14 ALL subtypes, generated from RNA-seq data from 1042 patients from five different ALL cohorts^3,28–30^. In a validation dataset, ALLCoP was able to reduce the false negative rate (FNR) from 8.95% to 3.5%. ALLCoP was then used to create prediction sets for 126 samples whose subtype was unknown at diagnosis. Finally, ALLCoP was validated on an additional two RNA-seq ALL subtype classifiers, ALLSorts^4^ and ALLCatchR^5^.

## METHODS

### Data

Publicly available gene expression (GEX) counts were obtained from five different studies ^3,28–30^ comprising RNA-seq data from a total of 1042 diagnostic ALL samples. The RNA-seq data were preprocessed, including the harmonization of subtype denotation and gene identifiers, batch correction with ComBat_seq and count normalization with TMM^31^. The preprocessed data were then subjected to ALL subtype prediction using ALLIUM^3^, a two-tiered, one vs. rest nearest shrunken centroid^32^ model. Model and data preprocessing details are in **Supplementary Methods**.

Samples whose known subtype was not recognized by ALLIUM and subtypes with n < 10 patients were excluded from the study. From the GSE227832 dataset^3^, the 207 samples that had been used to train the ALLIUM model were excluded. Of the 1042 samples meeting inclusion criteria, samples whose subtype was denoted as unknown in the original studies (n=126) and samples with multiple known subtypes (n=65) were analyzed separately, leaving 851 samples for ALLCoP calibration and validation.

### ALLCoP architecture

The conformal predictor in the present study, ALLCoP, was implemented using ALLIUM classifier outputs as inputs, for both the calibration process and subsequent formation of prediction sets (**Supplementary Figure S1**). We used *split conformal prediction*, which requires calibration and validation using a dataset independent from the training dataset of the underlying model^19^, and *conformal risk control*, which allows the application of CP to datasets with multiple true classes^33^. The latter produces prediction sets at a user-defined tolerance on the FNR, denoted α, and an FNR-controlling softmax threshold value*, lamhat*, above which classes are included in the prediction sets (**Supplementary Methods; Supplementary Figure S2**).

### ALLCoP validation

Using the ALLIUM predictions for all samples with a single known subtype (n=851), a series of CP cross-validation experiments were configured whereby each experiment consisted of multiple runs and in each run, the prediction dataset was shuffled and split: 90% of predictions were used for calibration of the conformal predictor using the conformal risk control^33^ algorithm, and 10% were used for validation.

First, we determined an optimal error rate (FNR tolerance) α which would result in the largest proportion of certain prediction sets (that is, prediction sets containing only one class). 1000 conformal prediction runs were initiated for each α value between 0.05 and 0.5, in increments of 0.05, and the sizes of the resulting prediction sets in the validation split were measured.

Second, we performed empirical validation to ensure that for a range of α values, the mean FNR was indeed no greater than the user-specified α. 1000 runs were executed with the optimal error rate α pre-selected above, and the FNR of every run (that is, the mean FNR of all the prediction sets in each run) was calculated and plotted. In addition, using 1000 runs per α value, the FNR of the ALLCoP prediction sets with α ∈ {0.1, 0.2, 0.3} were compared against the FNR of the uncalibrated ALLIUM classifier outputs, with the uncalibrated classifier outputs defined as a set containing the single top-scoring subtype.

Finally, we evaluated the mean FNR and mean set size stratified by ALL subtype, by running cross-validation experiments of ten thousand runs per α value in α ∈ {0. 075, 0. 1, 0. 15}.

### Prediction with ALLCoP

ALLCoP was re-calibrated in different configurations in order to generate prediction sets for subsets of the data. ALLCoP was calibrated on the samples with a single known subtype from the St. Jude Cloud dataset^28^ (n=594) at three different confidence levels α ∈ {0. 075, 0. 1, 0. 15} and used to generate prediction sets for the remaining n=257 samples with a single known subtype, as well as 65 samples with multiple known subtypes from the other three cohorts ^3,29,30^.

We then re-calibrated three instances of ALLCoP using all cases with single known subtype (n=851) at rates α ∈ {0. 075, 0. 1, 0. 15} and used them to generate prediction sets for the unknown-subtype cases (n=126).

### Code and data availability

All code and data used in this study are publicly available. Data was preprocessed for ALLIUM with allium_prepro^31^ and subjected to prediction with ALLIUM^34^. Conformal risk control was implemented as a Python package, conformist^35^, which was in turn used in ALLCoP^36^.

The St. Jude Cloud^28^ GEX counts and phenotypes are available from https://www.stjude.cloud/ under the accessions SJC-DS-1001 and SJC-DS-1009. The remaining GEX counts are available on the Gene Expression Omnibus (GEO): Diedrich et al./St. Jude Total Therapy XVI^29^ at accession number GSE161501; Krali, Heinäniemi et al.^3^ at GSE227832 and GSE228632; Tran et al.^30^ at GSE181157.

## RESULTS

### Uncertainty in ALL subtype classification

Our aim was to investigate if applying CP could quantify the uncertainty, and reduce the error rate, of RNA-seq based classifiers for determining molecular subtype of ALL. We applied ALLIUM^3^ to 1042 orthogonal RNA-seq samples from ALL patients from five studies ^3,28–30^ (**Supplementary Table S1**). The distribution of the molecular known subtypes from the five studies is shown in **Figure 1A**.

**Figure 1.**
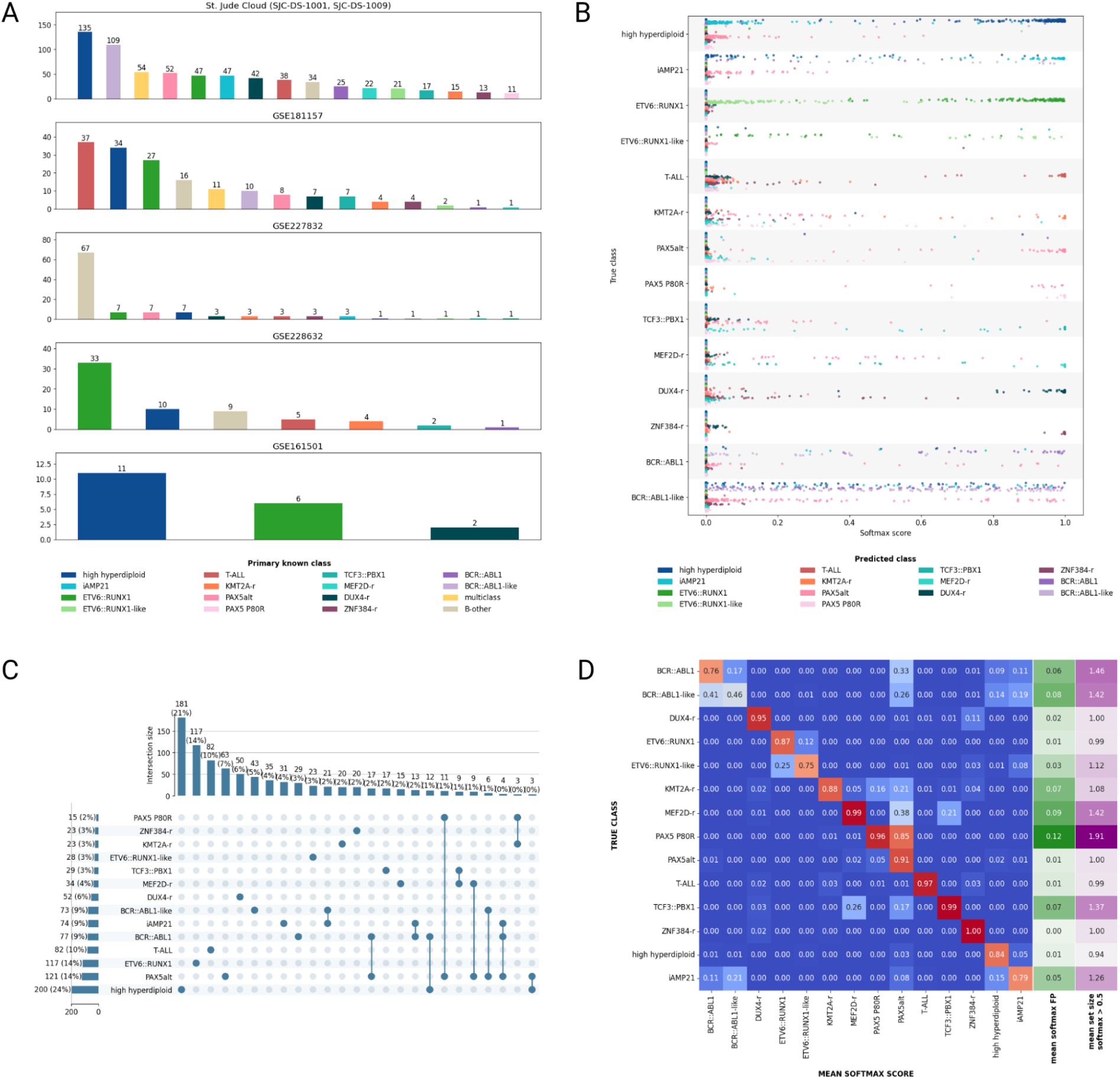
RNA-sequencing datasets and ALLIUM predictions used as input for ALLCoP. A) The subtype distributions for all 1042 samples used in the study. The remaining panels visualize ALLIUM predictions for the 851 samples with a single known subtype, including B) all softmax scores output by the model, stratified by true subtype and colored by predicted subtype C) the class membership of ALLIUM prediction sets, formed using a softmax threshold of 0.5 D) Heatmap mapping each true subtype to the mean softmax score per predicted subtype, with the green column showing the mean softmax scores of false positives and the purple column showing the mean size of the prediction sets with softmax threshold=0.5.

For the 851 samples with a single known subtype, the softmax scores of the ALLIUM predictions were unequally distributed by true subtype, with e.g. T-ALL and *ZNF384*-r showing consistently high scores for the correct subtypes only, while other subtypes such as *BCR::ABL1* and *BCR::ABL1-like* receiving high scores in more than one class (**Figure 1B**).

Using a softmax threshold of 0.5, we formed prediction sets comprised of all subtypes whose ALLIUM softmax score surpassed this value. 691 of the samples (81.2%) resulted in single-class prediction sets, with the most frequent sets containing a single prediction of high hyperdiploid (n=181) or *ETV6::RUNX1* (n=117). However, some subtypes were consistently found to co-occur with others, such as PAX5 P80R, where 78.6% of prediction sets containing this class also contained the subtype *PAX5*alt (**Figure 1C**).

Indeed, both subtypes PAX5 P80R and *PAX5*alt are defined by alterations of the *PAX5* gene, and using softmax scores alone ALLIUM has difficulty distinguishing between the two, although the true subtype typically receives a higher softmax score, with true PAX5 P80R receiving a mean softmax score of 0.96 versus 0.85 for *PAX5*alt. Again, using a softmax score cut-off of 0.5, the mean prediction set size, meaning the number of classes predicted per individual sample, varied depending on subtype, with the highest mean set sizes observed for the PAX5 P80R, *BCR::ABL1, BCR::ABL1*-like, and *MEF2D*-r subtypes (**Figure 1D**).

### Empirical error rate selection and False Negative Rate validation

Next, we developed and applied an ALL conformal predictor (ALLCoP) to the ALLIUM softmax scores (predictions), in order to 1) evaluate error reduction to predefined levels and 2) enable reporting of multiple potentially true subtype calls^37^.

Using ALLCoP, we created prediction sets using a predefined set of α levels (range 0.05 - 0.50), and a corresponding FNR-controlling softmax threshold value, lamhat, above which classes are included in the output prediction sets (**Figure 2A**; **Supplementary Table S2**).

**Figure 2.**
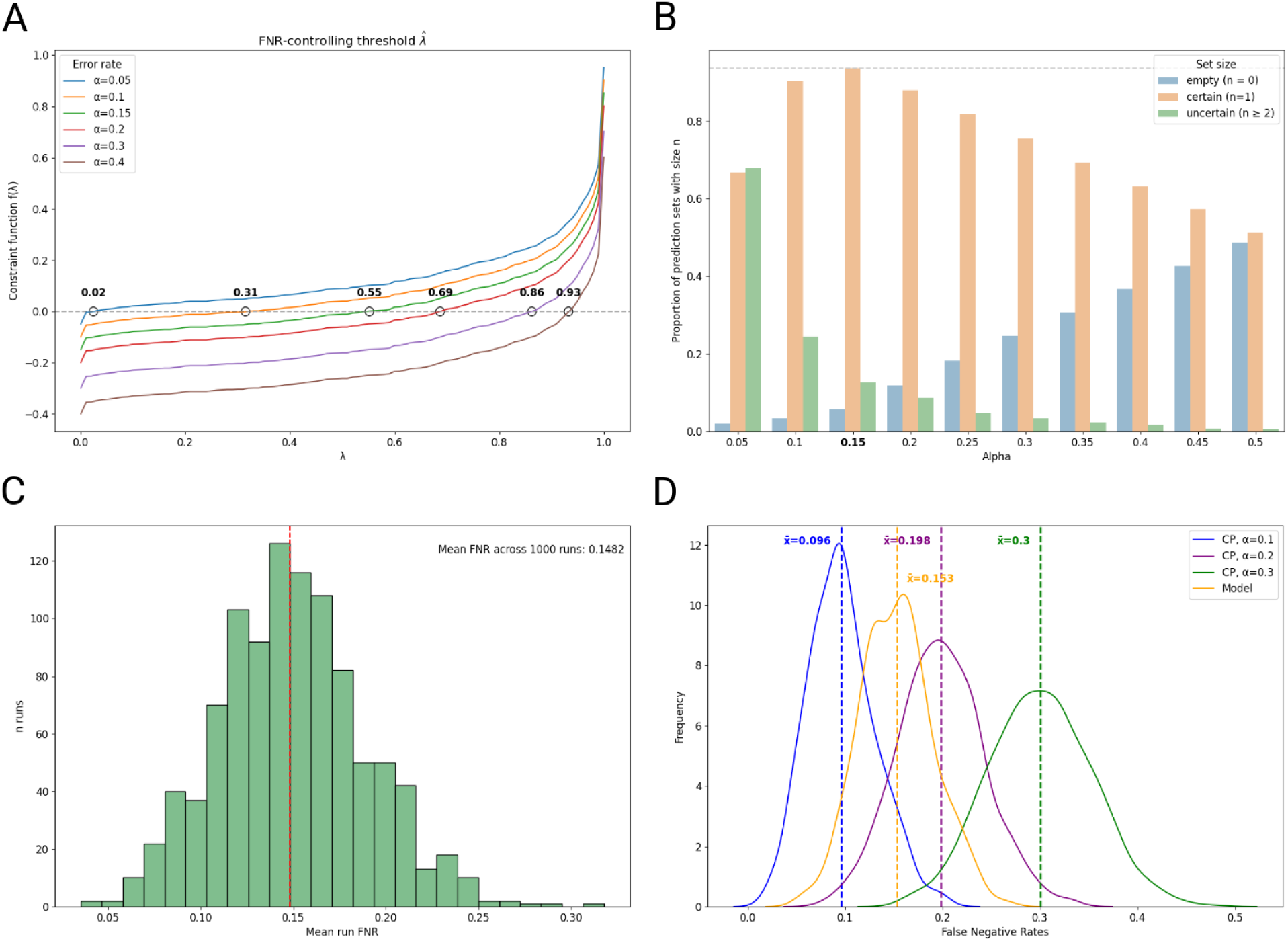
Empirical ALLCoP error rate selection and False Negative Rate (FNR) validation for the ALLIUM classifier. A) The output of the constraint function that selects the FNR-controlling model softmax threshold *lamhat* for a range of error rate values *α*. B) From an experiment of 1000 ALLCoP runs for a range of α values, pictured are the proportions of resulting prediction sets that were either empty, certain (size = 1) or uncertain (size >= 2). The gray dashed line and bolded number indicate the α value at which the highest proportion of certain prediction sets occurs. C) The mean FNRs of prediction sets produced in an experiment of 1000 ALLCoP runs with error rate α=0.15, empirically testing the overall FNR. D) The FNRs of the ALLCoP prediction sets produced at a range of α values, versus the FNR of the uncalibrated ALLIUM classifier outputs, defined as a set containing the single top-scoring subtype, in yellow.

Setting the FNR tolerance to α=0.15 produced the highest proportion of single-class prediction sets (81.6%). These were the most certain sets. This α also achieved a relatively small proportion of set sizes that either represented no prediction (5.7% empty sets) or uncertain prediction (12.7% of sets with size ≥ 2; **Figure 2B**). The mean FNR of the prediction sets was 14.82%, in line with the selected value of α=0.15 (**Figure 2C**).

Finally, in experiments of one thousand runs per α value, the FNR of ALLCoP prediction sets with α ∈ {0. 1, 0. 2, 0. 3} were compared against the FNR of the uncalibrated ALLIUM classifier outputs. In line with the conformal statistical guarantee, the FNRs of the ALLCoP prediction sets for ALLIUM were 9.6%, 19.8%, and 30.0%, respectively, while the mean FNR for the ALLIUM outputs was 15.3% (**Figure 2D**). FNRs for ALLCoP prediction sets at all three error rates were at or below the corresponding α values, indicating that the ALLCoP statistical guarantee holds across a diverse range of conditions, and regardless of the underlying model’s performance.

### ALLCoP performance by ALL subtype

Next, we evaluated ALLCoP performance by true ALL subtype. In order to obtain generalizations across the entire dataset, we ran a cross-validation experiment of ten thousand runs per α value with α ∈ {0. 075, 0. 1, 0. 15} using predictions from the 851 single-subtype samples, and evaluated the FNR and set sizes of the resulting prediction sets, *stratified* by subtype.

At α=0.075, the FNRs for the different subtypes ranged from 0-28% (**Figure 3A**), while the set sizes ranged from 1.0-2.50 (**Figure 3B**). At α=0.1, the FNRs ranged from 0-38% (**Figure 3C**) and set sizes from 0.98-2.0 (**Figure 3D**). At α=0.15, the FNRs ranged from 0-55% (**Figure 3E**) and set sizes from 0.93-1.91 (**Figure 3F**).

**Figure 3.**
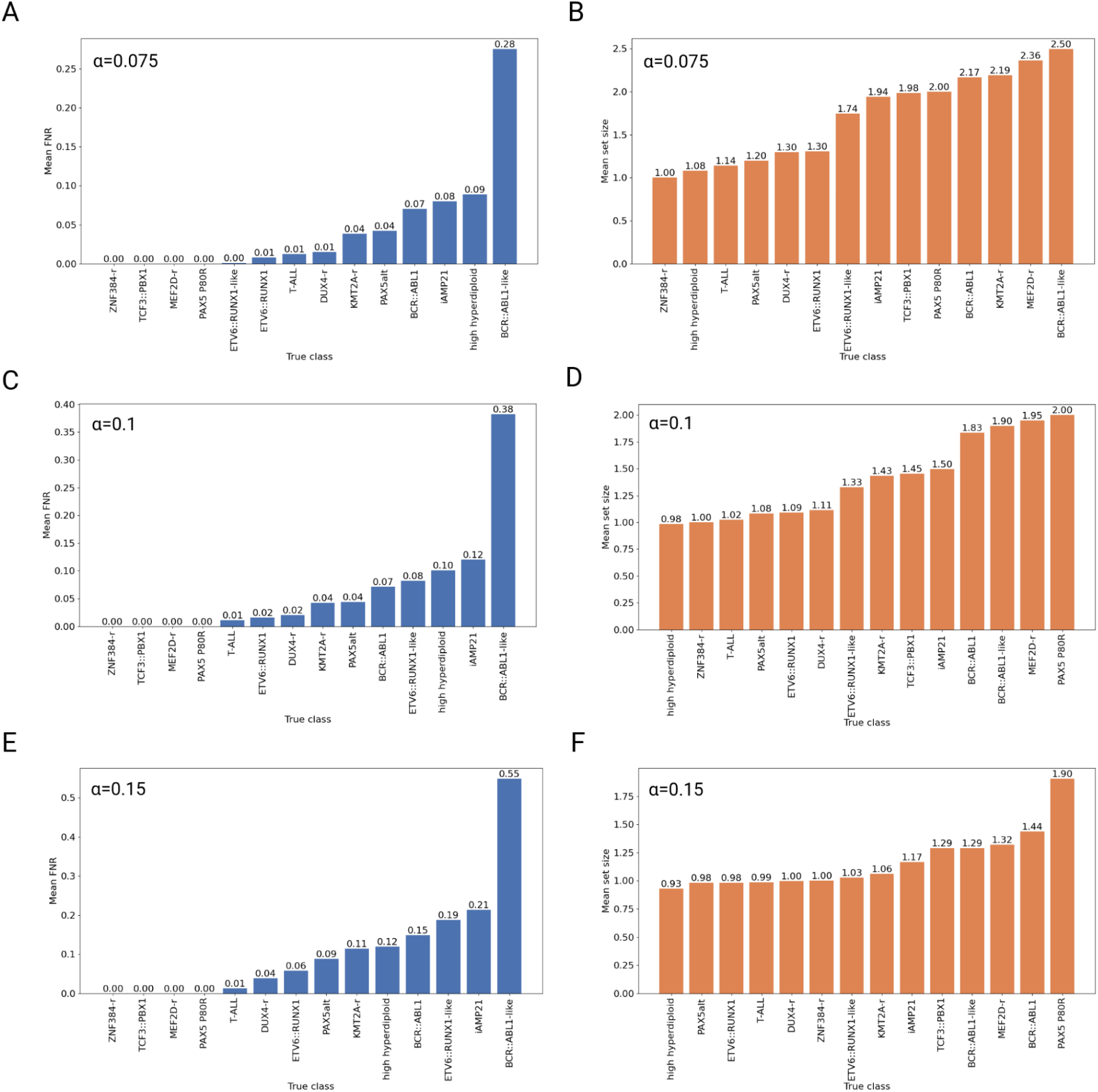
ALLCoP prediction sets generated from ALLIUM predictions. A) Mean False Negative Rates (FNRs) for prediction sets, α=0.075 B) Mean sizes for prediction sets, α=0.075 C) Mean FNRs for prediction sets, α=0.1 D) Mean sizes for prediction sets, α=0.1 E) Mean FNRs for prediction sets, α=0.15 F) Mean sizes for prediction sets, α=0.15

For subtypes *ZNF384*-r, *TCF3::PBX1* and *MEF2D*-r, the FNR remained 0% across all three α values, indicating high classifier certainty, and that the correct class was always included in the respective prediction sets. *BCR::ABL1*-like had the highest FNR across the three α values, from 28% at α=0.075 to 55% at α=0.15, and the highest set size of all subtypes at α=0.075 (2.50), indicating that subtypes with high uncertainty are increasingly included in prediction sets at lower α values, at the cost of larger prediction sets.

### Implicit error reduction in a validation dataset

After cross-validating the conformal guarantee in ALLCoP and assessing its performance across ALL subtypes, we then re-calibrated individual instances of ALLCoP to produce prediction sets for three data subsets: first, for a validation subset of the data with single known subtype; second, for samples with multiple known subtypes; and finally, for samples of unknown subtype. We aimed to show that the conformal predictor can be used to reduce error in validation sets where subtype is known, and to produce fewer empty predictions for samples with previously unknown subtype.

Using ALLIUM predictions from the St. Jude Cloud samples with single known subtype (n=594)^28^, we recalibrated ALLCoP with α ∈ {0. 075, 0. 1, 0. 15} and used it to obtain prediction sets on a validation set of the remaining samples with a single known subtype (n=257)^3,29,30^ (**Supplementary Table S3**).

In this validation set, 234 ALLIUM predictions were correct, 19 were wrong, and 4 were empty. Of these 23 samples with incorrect or empty predictions, ALLCoP prediction sets contained the correct class for six cases at α=0.15, 13 cases at α=0.1 and 14 cases at α =0.075. Overall, in this validation set, the uncalibrated ALLIUM model had an FNR of 8.95%, which was reduced to 6.61% at α=0.15, 3.89% at α=0.10 and 3.50% at α=0.075; the trade-off was an increasing mean set size: 1.11 at α=0.15, 1.31 at α=0.10 and 1.53 at α=0.075 (**Supplementary Table S4**). A comparison between ALLIUM class predictions, ALLCoP sets, and sets containing classes where the softmax threshold was greater than 1-α is shown for α=0.075 (**Figure 4A**), α=0.1 (**Figure 4B**), and α=0.15 (**Figure 4C**). The mean FNR and mean set size stratified by subtype is tabulated in **Supplementary Table S5**. Notably, the FNR of the *BCR::ABL1*-like subtype, which was 90.9% in the uncalibrated ALLIUM output, was reduced to 36.36% at α=0.075. Also of note, the PAX5alt subtype frequently co-occurred with other subtypes, appearing in 10 of 29 multi-class prediction sets at α=0.15 (34.48%), 20 of 65 at α=0.10 (30.77%), and 31 of 103 (30.10%) at α=0.075.

**Figure 4.**
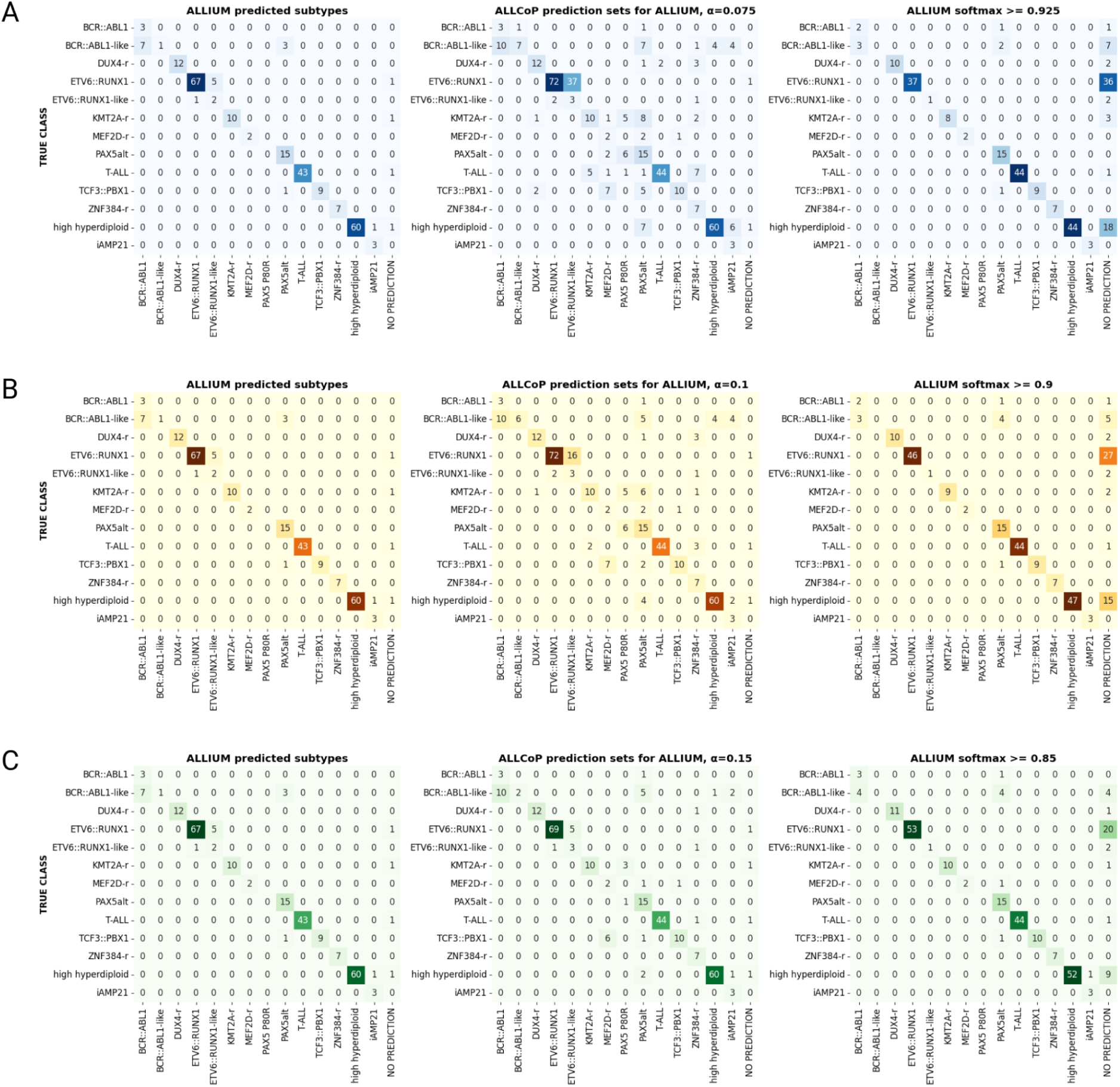
The concordance between ALLIUM single-class point predictions, ALLCoP sets, and sets of classes where the softmax threshold was greater than 1-α in the validation dataset with single known subtype, at pre-selected false negative rates of A) α=0.075, B) α=0.1 and C) α=0.15.

We then used the same conformal predictors to obtain prediction sets on a validation set of the samples with multiple known subtypes (n=65)^28,30^ (**Supplementary Table S6**). Briefly, the uncalibrated ALLIUM model had an FNR of 61.54%, which was reduced to 23.85% at α=0.075 (**Supplementary Table S7**; performance by subtype in **Supplementary Table S8**). The prediction sets are visualized in **Supplementary Figure S3**.

### Prediction sets for unknown subtype cases

We collected the cases with unknown subtype from all studies (n=126), calibrated three instances of ALLCoP using all cases with single known subtype (n=851) at error rates α ∈ {0. 075, 0. 1, 0. 15}, and used them to generate prediction sets for the unknown-subtype cases (**Supplementary Table S9**).

Using the uncalibrated model, ALLIUM issued predictions for 97 cases at the model default softmax cutoff of 0.5, leaving 29 empty predictions (23%). At α=0.15, 34 of the prediction sets were empty (27%) (**Figure 5A**), but at α=0.10, this number dropped to 26 empty sets (21%; **Figure 5B**), and at α=0.075, only 21 of the prediction sets were empty (17%; **Figure 5C**). Across all three error rates, the most commonly predicted subtypes were *PAX5*alt, *DUX4*-r, and iAMP21.

**Figure 5.**
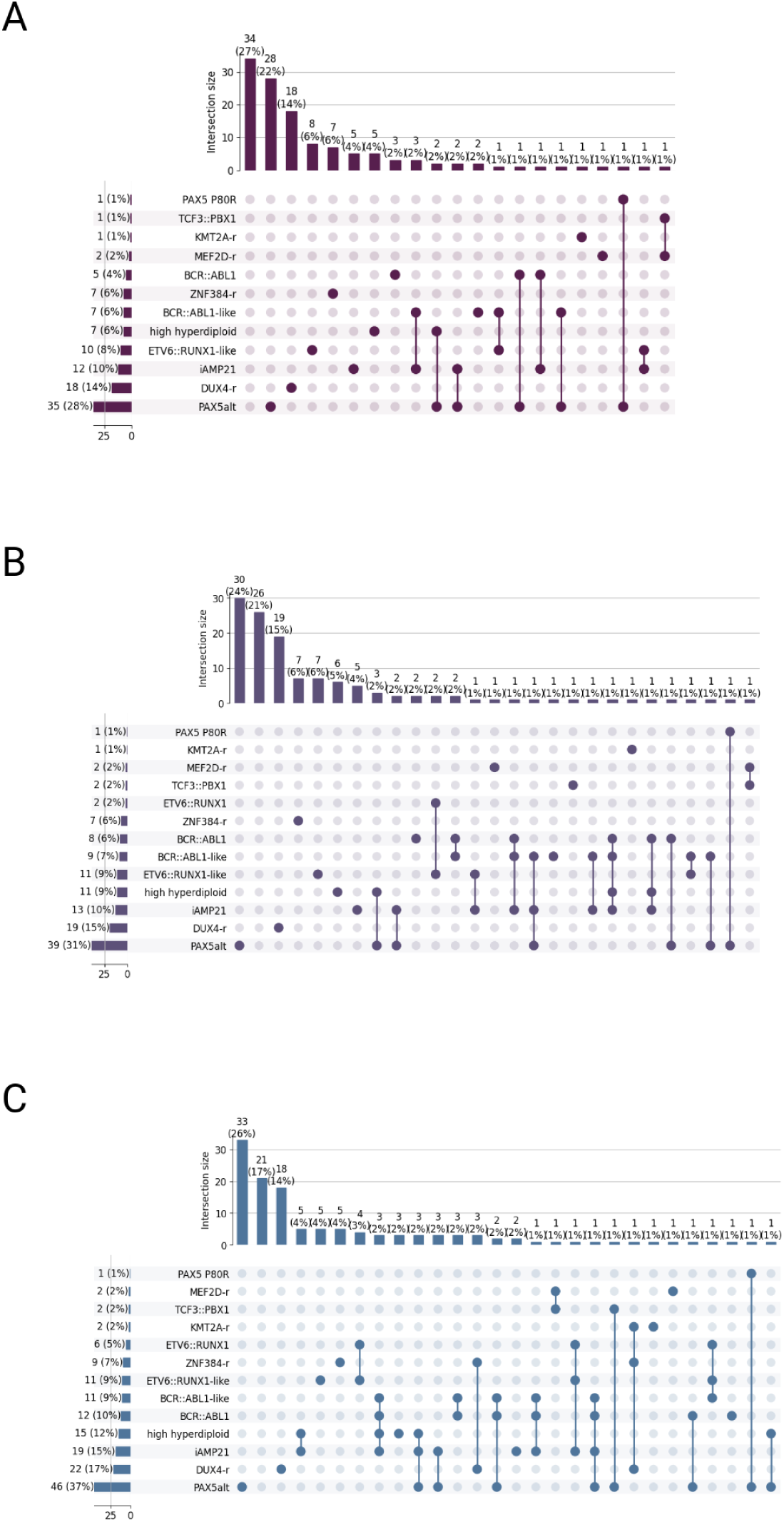
ALLCoP prediction sets for 126 previously unclassified B-ALL cases using predictions from ALLIUM. The upset plots represent class membership counts, with unattached dots representing single-class prediction sets and connected dots representing multi-class sets. Visualized are prediction sets for pre-selected False Negative Rates of A) α=0.15, resulting in 34 empty sets, 78 certain sets, and 14 uncertain sets B) α=0.10, resulting in 26 empty sets, 80 certain sets, and 20 uncertain sets C) α=0.075, resulting in 21 empty sets, 69 certain sets, and 36 uncertain sets.

### ALLCoP validation on additional RNA-seq ALL subtype classifiers

ALLIUM is one of numerous RNA-seq classifiers developed for molecular subtype determination in ALL^4–6^. To evaluate the generalizability of ALLCoP to other classifiers, we selected two of the other models, ALLCatchR^5^ and ALLSorts^4^ and generated predictions for RNA-seq data from samples that were not used for training these models^3,29^, (**Supplementary Figure S4; Supplementary Tables S10-S11**). The softmax scores generated by ALLCatchR resulted in discrete distributions (**Figure 6A**). The subtypes with most frequent uncertainty were iAMP21 (high softmax scores for *BCR::ABL1*-like, *ETV6::RUNX1*-like, and high hyperdiploid) and *PAX5*alt (high softmax scores for *BCR::ABL1*-like). Similar to ALLCatchR, ALLSorts was most uncertain for iAMP21, often confusing it for the same classes as ALLCatchR (**Figure 6B**). ALLCatchR generated overwhelmingly certain predictions, with the mean prediction set size (using a softmax cutoff of 0.5) never surpassing 1.0 and remaining over 0.99 for all classes except *PAX5*alt (0.93) and iAMP21 (0.69; **Figure 6C**). Similarly, ALLSorts had a mean set size of over 0.90 for all classes except *KMT2A*-r (0.83) and iAMP21 (0.50), although it had more uncertain sets, with a mean set size >1.0 for *BCR::ABL1*-like (1.14), *ZNR384*-r (1.12) and *PAX5*alt (1.07; **Figure 6D**). In ALLSorts, as with ALLIUM prediction sets, *PAX5*alt was the subtype most frequently observed in multiclass predictions, although in the discrete model outputs of ALLCatchR, this was not observed.

**Figure 6.**
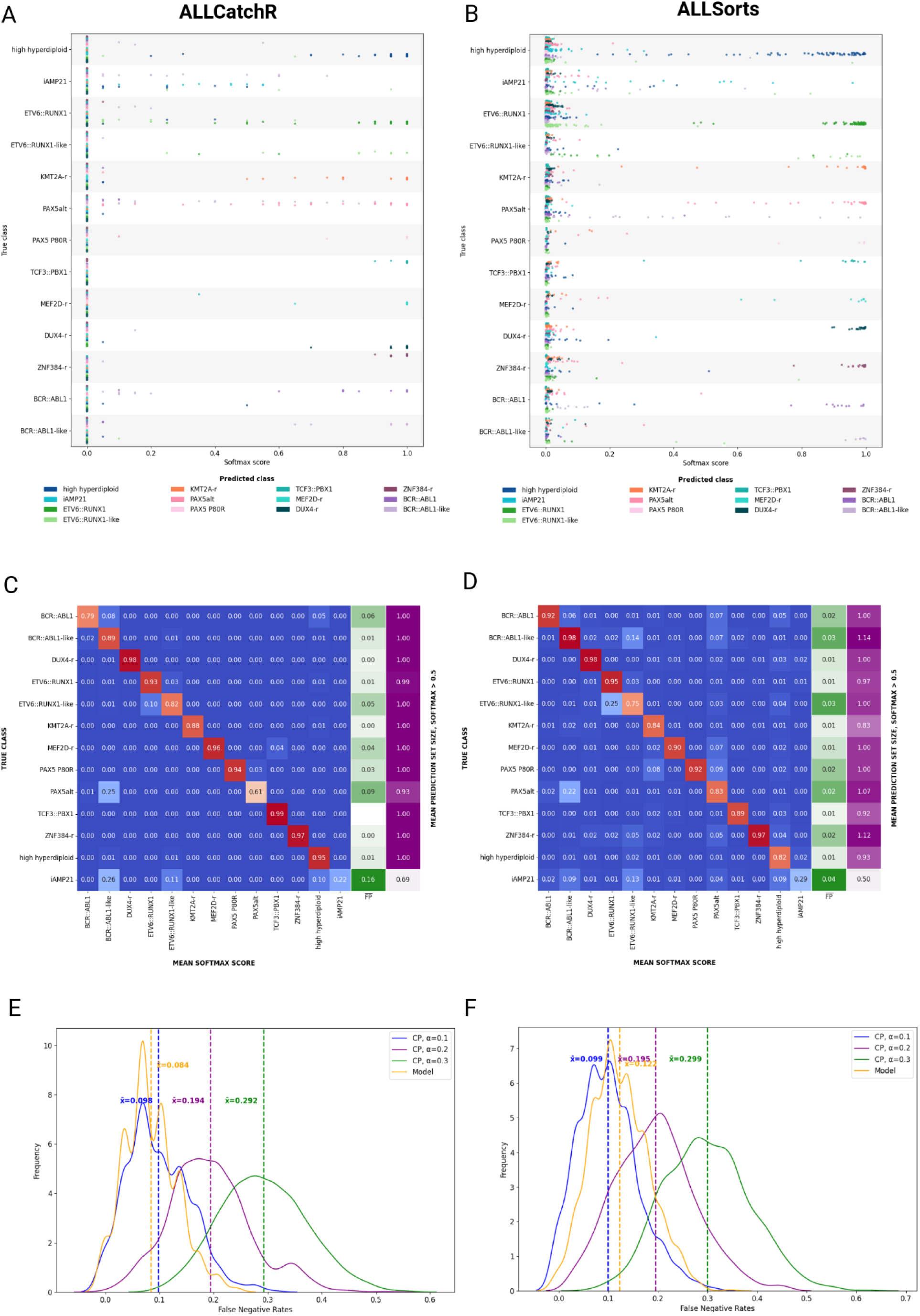
Softmax predictions and ALLCoP results for two ALL RNA-seq classifiers, ALLCatchR and ALLSorts. All softmax scores output by the model, stratified by true subtype and colored by predicted subtype, for A) ALLCatchR and B) ALLSorts. Heatmaps mapping each true subtype to the mean softmax score per predicted subtype, with the green column showing the mean softmax scores of false positives and the purple column showing the mean size of the prediction sets with softmax threshold=0.5, for C) ALLCatchR and D) ALLSorts. False Negative Rate of ALLCoP prediction sets vs. uncalibrated model outputs for E) ALLCatchR and F) ALLSorts, with the uncalibrated model outputs in yellow.

Next, we applied ALLCoP to the softmax scores and evaluated the output (**Supplementary Tables S12-S13**). Similarly to ALLIUM, ALLCoP produced prediction sets at defined FNR for AllCatchR and ALLSorts (**Figure 6E-F**). Empirical error rate selection was performed for the two classifiers, showing an optimal error rate of α=0.1 for ALLCatchR (softmax threshold of 0.44) and α=0.05 for ALLSorts (softmax threshold of 0.18; **Supplementary Figure S5**). All together, these results further demonstrate the robustness and generalizability of ALLCoP for error reduction in RNA-seq ALL classifiers.

## DISCUSSION

In this study, we present the first application of conformal risk control to RNA-seq-based ML classification of ALL. CP provides several key advantages that enhance the robustness and interpretability of ML models in this context. These include statistical performance certification for predictions, the ability to guarantee a user-defined error rate irrespective of the model’s intrinsic performance, and explicit quantification of prediction uncertainty, with larger prediction sets in cases of higher uncertainty. Moreover, CP inherently facilitates implicit error reduction, contributing to more reliable classification outcomes.

Recent and forthcoming legislation, such as the EU AI Act, emphasizes regulatory requirements including transparency, explainability, and accountability in AI systems^38,39^ In this context, the quantification of model robustness and confidence are essential^40^. This necessitates methodologies that go beyond the uncalibrated softmax outputs typically generated by classifiers, as these outputs lack a direct connection to empirical probabilities^41,42^.

In this study, we observed the properties of CP when applied to ALL subtype prediction. One important observation was that the mean error rates of the conformal prediction sets varied significantly by subtype. For example, for *ZNF384*-r we observed a FNR of 0% across all α values and classifiers, despite the fact that this subtype had among the fewest calibration examples (n=24 ALLIUM predictions, n=17 ALLCatchR/ALLSorts predictions). This suggests that this subtype has highly distinctive biological signals that can be reliably detected in transcriptomic data to differentiate it from the other ALL subtypes.

In contrast, the *BCR::ABL1*-like subtype had the highest mean FNR (36-82%) across the error rates in ALLIUM ALLCoP prediction sets (although in the other classifiers it ranged from only 0-4%). The subtype *BCR::ABL1*-like was established to define cases that share a similar transcriptomic profile with *BCR::ABL1*, but lack this canonical subtype-defining fusion gene^43^, and in ALLIUM, which was only trained on n=7 *BCR::ABL1*-like samples, the model has trouble differentiating between the two subtypes. Further, there are inconsistencies in the definition of “ground truth” for this subtype across the different ALL studies and cohorts included herein. Some cohorts rely on gene expression clustering to classify cases^30^, whereas others use genetic aberrations of JAK-STAT pathway genes and/or the overexpression of *CRLF2* as subtype-defining critera^8,44^. These discrepancies highlight a limitation of the current study, as variability in subtype definitions may violate the requirement for data exchangeability in CP^19^ and may have thus reduced its precision. In future studies, this shortcoming can be addressed by leveraging CP techniques that are more robust under distributional drift and that circumvent this need for data exchangeability^45^. In addition, calibrating a conformal predictor to a user-specified error rate not just over the entire dataset, but within individual classes, can help overcome class-specific performance inequities^46^.

The variability of CP set size, referred to as *set adaptivity*, serves as a valuable metric of uncertainty. This reflects both the performance of the classifier and the irreducible uncertainty arising from the inherent complexity of the features being classified. In our study, the *PAX5*alt subtype most frequently appeared in multi-class prediction sets generated using ALLIUM^3^ and ALLSorts^4^. Given that alterations in the *PAX5* gene are observed in over one-third of BCP-ALL patients, as reported in prior studies^37,47^, the multi-class conformal prediction sets containing *PAX5*alt may not solely indicate classifier uncertainty, but could also point to biologically relevant secondary aberrations with potential clinical importance.

Finally, we explored the concept of implicit error reduction within the framework of ALL RNA-seq classifiers. In the validation cohorts, we found that lowering the α value resulted in the expected reduction in the mean FNR of the prediction sets. Likewise, for samples with unknown subtypes, a lower α value resulted in fewer empty prediction sets, demonstrating improved model utility.

However, this approach involves a tradeoff: reducing the α value increases the likelihood of including the true class in the prediction sets but results in larger and less precise sets^48^. The ability to select an α value tailored to specific requirements offers significant flexibility. For instance, a higher tolerance for error may be acceptable in research contexts where exploratory analysis is prioritized. Conversely, high-stakes applications demand stricter error rates to ensure reliable and actionable predictions ^49,50^. This adaptability underscores the potential of CP as a versatile tool for integrating ML into both research and clinical workflows, balancing precision with reliability based on user-defined thresholds.

In summary, this study demonstrates the potential of CP to enhance the robustness, transparency, and adaptability of RNA-seq-based machine learning classifiers for ALL subtyping. By addressing both predictive uncertainty and error management, our findings pave the way for integrating advanced AI methodologies into clinical workflows while aligning with emerging regulatory requirements.

## ACKNOWLEDGMENTS

This study was conducted with support from the Swedish Childhood Cancer Foundation (MT2022-0006 and HFT2023-0011), the Swedish Research Council (#2019-01976), and the Göran Gustafsson Foundation. We thank the SciLifeLab National Genomics Infrastructure (NGI), SNP&SEQ Technology Platform, for assistance with generating RNA-seq data. NGI is funded by SciLifeLab, the Swedish Research Council, and the Knut and Alice Wallenberg Foundation. The computations and data handling were enabled by resources provided by the National Academic Infrastructure for Supercomputing in Sweden (NAISS), partially funded by the Swedish Research Council through grant agreement no. 2022-06725. Special thanks to Kim Kultima for the constructive feedback on this manuscript.

## Notes

### Competing Interest Statement

The authors have declared no competing interest.

