## Supplementary Material for "Error reduction in leukemia machine learning classification with conformal prediction"

---

Mariya Lysenkova Wiklander, Dave Zachariah, Olga Krali, Jessica Nordlund

### Supplementary Information

Supplementary Methods
- RNA-seq ALL subtyping classifiers
- Conformal risk control
- ALLIUM data preprocessing and filtering
- ALLSorts and ALLCatchR data preprocessing and filtering
- Reference genome
- Conformist: a Python package for conformal prediction

Supplementary Figures
Supplementary References

### Supplementary Methods

#### RNA-seq ALL subtyping classifiers

Three RNA-seq classifiers were used in the present study: ALLIUM[1], ALLSorts[2] and ALLCatchR[3].

ALLIUM is a nearest shrunken centroid[4] model that enables the molecular classification of ALL cases into 17 known subtypes. It was designed as a two-tiered, one vs. rest model. ALLIUM was trained on 12 ALL groups, plus a control group, where each group contains one or more subtypes. For instance, the aneuploidy group contains the hyperdiploid, low hyperdiploid and iAMP21 subtypes, while the *ETV6* group contains *ETV6::RUNX1* and *ETV6::RUNX1*-like. The optimization process resulted in the discovery of transcriptomic signatures not only specific to groups, but also to subtypes vis-a-vis other, similar subtypes. For example, signatures were detected that could differentiate specifically between *ETV6::RUNX1* and *ETV6::RUNX1*-like, or between the different aneuploidy types. Attempts were made to re-train the model using a "flat" structure, using no grouping of subtypes; however, predictions on a number of test data points resulted in softmax scores close to 1.0 for multiple, biologically similar classes. Therefore the decision was made to retain the two-tiered structure, which has potential implications for the distribution of softmax scores in the present study.

ALLSorts is also a hierarchical model, using a set of logistic regression classifiers to classify 18 known subtypes, while ALLCatchR is a hybrid system using linear support vector machines and nearest-neighbor models to predict 21 ALL subtypes.

For the purposes of the present study, we use only the final subtype predictions in calibrating the conformal predictor, treating any intermediate scores, such as the initial group level predictions in ALLIUM and ALLSorts, as part of the "black box" of the machine learning model.

#### Conformal risk control

The coverage guarantee of conformal risk control states that, for an input *x* with a user-specified error rate of α=0.1, the prediction set *y* contains an average of at least 90% of the true classes[5]. The following from Angelopoulos et al. is used to implement this guarantee:

$$\hat{\lambda} = \inf\left\{\lambda : \frac{1}{n+1}\sum_{i=1}^{n} s(X_i, Y_i) > \lambda + \frac{1}{n+1} \leq \alpha\right\} = \underbrace{\inf\left\{\lambda : \frac{1}{n}\sum_{i=1}^{n} s(X_i, Y_i) \leq \lambda \geq \frac{\lceil (n+1)(1-\alpha)\rceil}{n}\right\}}_{\text{conformal prediction algorithm}}.$$

The selection of a suitable error rate is a tradeoff between the inherent desirability of a lower False Negative Rate (FNR), and a crucial limitation of conformal prediction (CP): as the FNR approaches zero, the prediction sets become too large to be meaningful. Here, α refers to the user-selected error rate, while λ refers to the softmax threshold value above which a class is included in the conformal prediction sets. A constraint function is applied to the calibration dataset in order to find the optimal FNR-controlling threshold $\hat{\lambda}$ so that the mean FNR for a prediction set is less than or equal to α. Prediction sets are then created so that:

$$\{prediction\ set\ for\ x\} = all\ classes\ with\ classifier\ softmax\ score \geq \hat{\lambda}.$$

#### ALLIUM data preprocessing and filtering

The primary and secondary subtypes specified per sample in the phenotype metadata file provided by St. Jude Cloud were standardized to match the subtype names used by ALLIUM. A number of St. Jude subtypes were merged in order to map to ALLIUM subtypes. Although the St. Jude dataset differentiates between several different T-ALL subtypes, ALLIUM supports a single umbrella T-ALL subtype; therefore, all T-ALL samples were assigned to this class. Likewise, the St. Jude classes *Ph-like_non CRLF2* and *Ph-like_CRLF2* were consolidated to *BCR::ABL1-like; KMT2A* and *KMT2A-like* were consolidated to *KMT2A*-r*; ZNF384* and *ZNF384-like* were consolidated to *ZNF384*-r.

Samples were filtered out where the primary subtype of the patient was either unknown, not recognized by ALLIUM, or not recognized by the International Consensus Classification (ICC) of ALL[6]. In addition to the samples with unknown subtype at diagnosis, samples with the following St. Jude subtypes were filtered out: *CRLF2(non-Ph-like), Dicentric, Down, IKZF1 N159Y, Low hyperdiploid, Low hypodiploid, Near haploid*, and *ZEB2/CEBPE*. A total of 87 samples were filtered out, with 648 samples remaining.

Samples with a secondary subtype unrecognized by ALLIUM were retained in the dataset, but their secondary subtype was removed. There were 17 such samples, with the secondary subtypes of *CRLF2(non-Ph-like), Dicentric, Down*, and *Low hyperdiploid*.

The data preprocessing code is available in the allium_prepro repository[7].

#### ALLSorts and ALLCatchR data preprocessing and filtering

ALLSorts and ALLCatchR were both trained using the St. Jude Cloud dataset, rendering this dataset unsuitable for use with these classifiers in the present study, as such usage would violate the split CP assumption of independence between model training and conformal calibration datasets[8]. Therefore, for these classifiers, we merged WTS and phenotype data from three external sample sets: GSE227832 (n=328), GSE228632 (n=65) and GSE161501 (n=19). After filtering, 292 samples remained across 13 known subtypes.

Because known subtypes for the data were denoted using ALLIUM subtype conventions, the class names from ALLSorts and ALLCatchR predictions were translated to the ALLIUM equivalents.

For ALLSorts, the *BCL2/MYC, HLF, IKZF1 N159Y, Low hypodiploid,* and *Near haploid* subtypes were not represented in the data and were therefore removed from the prediction matrix. For ALLCatchR predictions, the following subtypes were removed from the prediction matrix, as they were not represented in the data: *BCL2/MYC, CDX2/UBTF, CEBP, HLF, IKZF1 N159Y, Near haploid*.

Neither ALLSorts nor ALLCatchR were trained to recognize *T-ALL*, so these cases were removed from their prediction matrices.

#### Reference genome

The reference genome used for this project was GRCh38.103. Gene symbols from all calibration and validation datasets were converted to Ensembl identifiers in accordance with this genome version. A Python utility called "Gene Thesaurus" (https://github.com/Molmed/gene-thesaurus) was implemented to leverage the HUGO Gene Nomenclature Committee database provided by EMBL-EBI in order to translate between gene symbols and Ensembl identifiers, as well as between different versions of gene names.

#### *Conformist*: a Python package for conformal prediction

ALLCoP was refactored into a Python package that can be used to calibrate conformal predictors and create the visualizations presented in the current study, for any classifier that outputs softmax scores. In addition to the conformal risk control predictor, *conformist* also offers a “vanilla” CP implementation that can be used for single-class classification problems. Further details are available in the README.md file for this project’s repository: https://github.com/Molmed/conformist.

#### Supplementary Figures

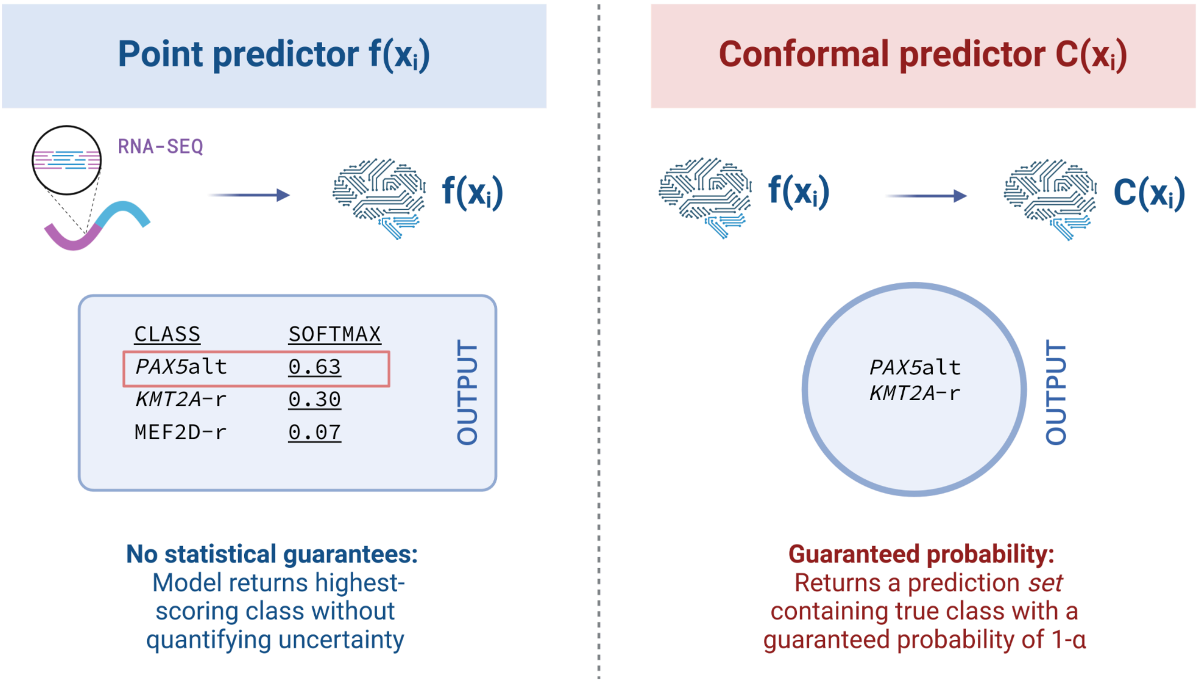


**Supplementary Figure S1**. Conformal prediction uses outputs from a machine learning model as inputs to calibrate a conformal predictor that is subsequently used to create statistically guaranteed prediction sets at a user-specified error level α. Created in https://BioRender.com

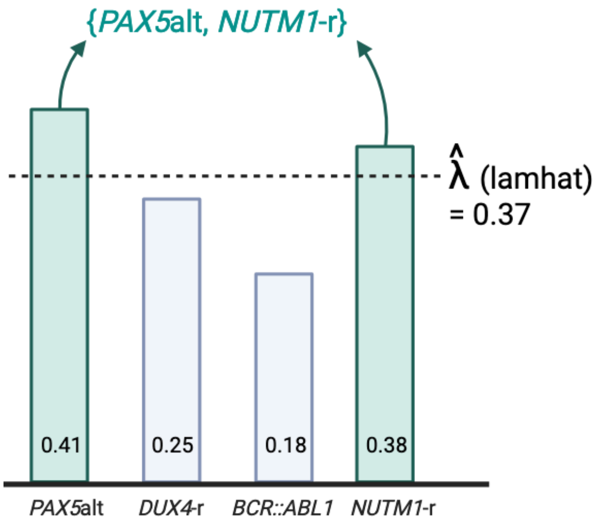


**Supplementary Figure S2**. A schematic showing how prediction sets are formed using the softmax threshold lamhat which is determined during the conformal calibration process. Created in https://BioRender.com

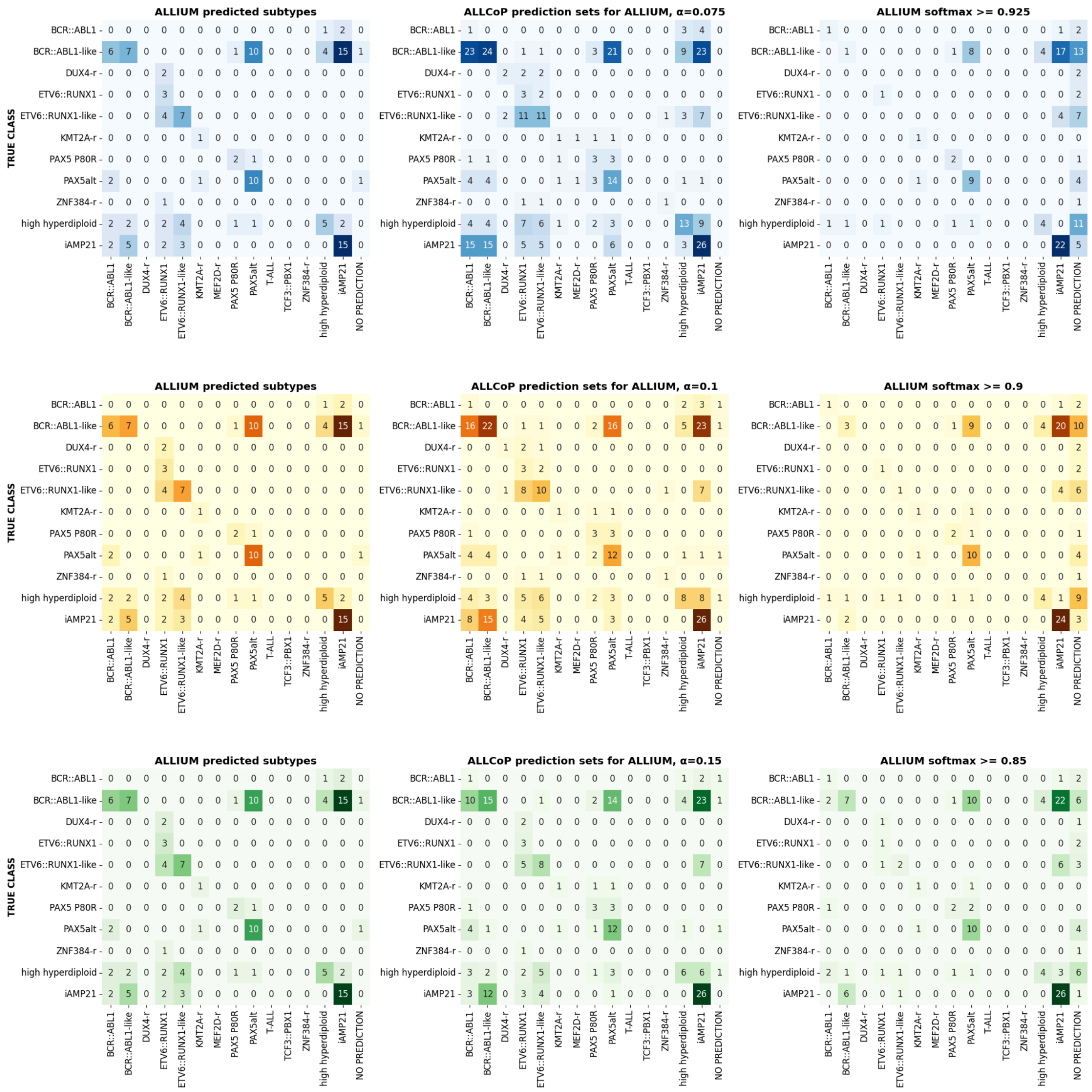


**Supplementary Figure S3. The concordance between ALLIUM single-class point predictions, ALLCoP sets, and sets of classes where the softmax threshold was greater than 1-α in the validation dataset with multiple known subtypes**, at pre-selected false negative rates of A) α=0.075, B) α=0.1 and C) α=0.15. In this validation set, ALLIUM predicted at least one correct class in 50 of the samples, 14 were wrong, and 1 was empty. Of these 15 incorrect or empty predictions, ALLCoP prediction sets contained the correct class for 6 cases at α=0.15, 12 cases at α=0.1 and 15 cases at α=0.075.

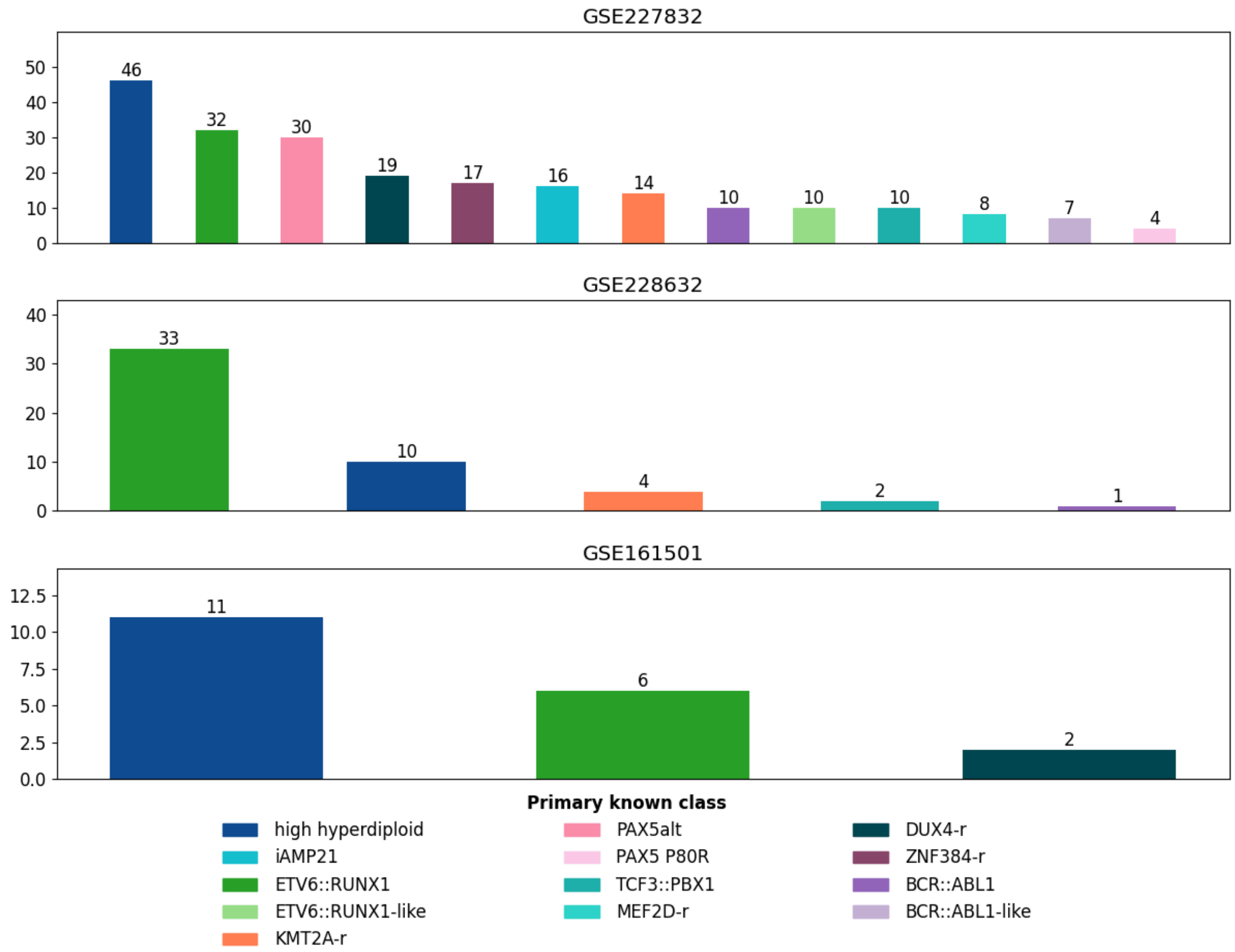


**Supplementary Figure S4. RNA-sequencing datasets used as input for ALLCatchR and ALLSorts.**

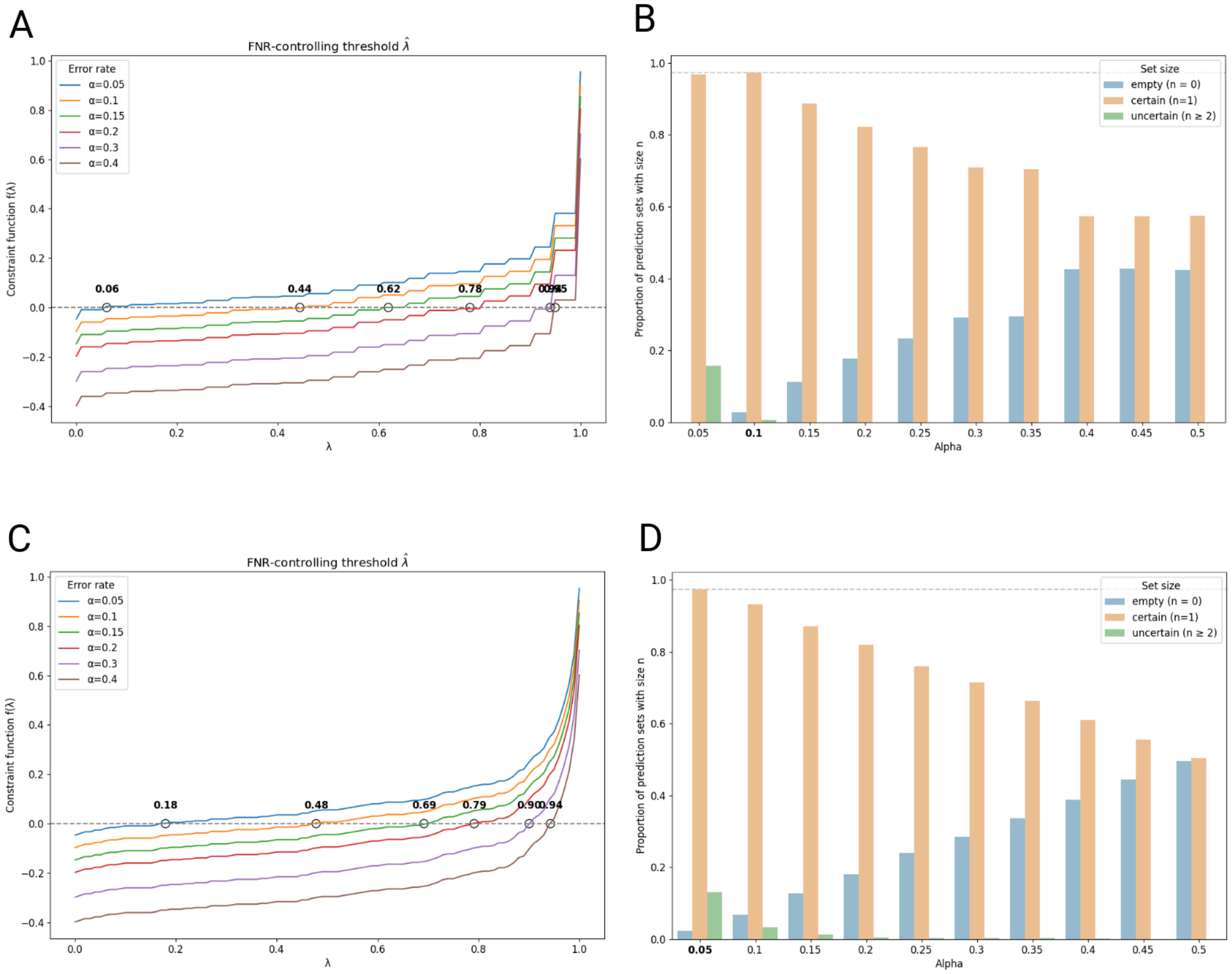


**Supplementary Figure S5. Empirical ALLCoP error rate and False Negative Rate-controlling softmax threshold selection for two ALL RNA-seq classifiers.** A) FNR-controlling softmax threshold selection for ALLCatchR B) Empirical error rate selection for ALLCatchR C) FNR-controlling softmax threshold selection for ALLSorts D) Empirical error rate selection for ALLSorts.
